# Comparative proteomic analysis of the ECM composition of the human omentum and mesentery, the main sites of ovarian cancer metastasis

**DOI:** 10.1101/2025.07.28.667199

**Authors:** James M. Considine, Clarissa Gomez, Dharma Pally, Ning Yang, Isra N. Taha, Jenna Sorenson, Erin G. Brooks, Pamela K. Kreeger, Alexandra Naba

**Author notes:** **Corresponding Authors: Pamela K. Kreeger,** University of Wisconsin School of Medicine and Public Health, Department of Pathology and Laboratory Medicine, 1111 Highland Ave, WIMR 5037, Madison, WI 53705; **Alexandra Naba**, University of Illinois Chicago, Department of Physiology and Biophysics, 1853 W. Polk Street, Suite CMWT 522, Chicago, IL 60612.

## Abstract

Due to its limited symptoms, high-grade serous ovarian cancer (HGSOC) has frequently metastasized extensively throughout the peritoneal cavity prior to its diagnosis, resulting in an overall five-year survival rate of less than 50%. The greater omentum and the small bowel mesentery are two of the most common metastatic sites in advanced HGSOC. However, the mechanisms underlying HGSOC metastatic tropism remain unknown. The extracellular matrix is a complex and dynamic meshwork of proteins that provides biochemical and mechanical signals to surrounding cells and has been shown to drive the dissemination of several cancer types to preferential distant sites. Here, using histological assessment and proteomics, we examined the composition of the extracellular matrix of paired omentum and mesentery samples from disease-free adult females. We found that the fibrillar collagen content of the mesothelial layer of the omentum was significantly higher than that of the mesentery. Using ECM-focused proteomics, we further defined the ECM composition – or matrisome – of these two tissues. We found that over 90% of the proteins detected were shared between the omentum and mesentery. Our analysis also revealed small subsets of tissue-specific ECM proteins. Future work will aim to test the possible functional contributions of these ECM proteins to HGSOC metastatic tropism. To facilitate the reuse of our dataset, we have deposited the raw mass spectrometry data and accompanying metadata files to the ProteomeXchange Consortium with the dataset identifier PXD061586.

## INTRODUCTION

Due to its limited symptoms, high-grade serous ovarian cancer (HGSOC) has frequently metastasized extensively throughout the peritoneal cavity prior to its diagnosis, resulting in an overall five-year survival rate of less than 50% ^1–4^. The greater omentum and the small bowel mesentery are two of the most common metastatic sites in advanced HGSOC ^5^. Nearly 80% of patients have a metastasis on the omentum, which is commonly removed by omentectomy during a debulking ^6^. In contrast, approximately 30% of patients present with tumors on the small bowel mesentery ^7^, with disease in this site primarily treated by chemotherapy. The difference in treatment approach stems from their unique anatomical roles. While both are components of the visceral peritoneum, double-layered tissues that cover and provide structure for the abdominal organs, the greater omentum is suspended loosely between the intestines and the peritoneal wall and has non-essential roles in fat storage and inflammation ^8^. In contrast, the mesentery surrounds the intestines and is heavily vascularized to provide the blood supply for these critical organs ^9^. Mesenteric involvement can prevent complete cytoreduction ^10,11^, which is associated with a worse outcome ^12^. Therefore, identifying the mechanisms that support metastasis to the mesentery could provide treatments that would improve HGSOC patient prognosis.

Due to its more common involvement and frequent removal during surgical debulking, the greater omentum has been extensively studied to identify factors that support HGSOC metastasis ^13–15^. Studies have demonstrated a loss of adipocytes as tumors progress in the omentum and identified mechanisms by which tumor cells utilize lipids stored in adipocytes for energy to support proliferation ^16^. Another common hypothesis for the involvement of the omentum in HGSOC is the presence of ‘milky spots’ ^17^, clusters of leukocytes and macrophages that may secrete factors to attract tumor cells ^18^ or alter the tissue microenvironment to support adhesion ^19^. However, neither of these observations would explain why the omentum is preferred to the mesentery, as both tissues have adipose and milky spots.

The extracellular matrix (ECM) is a complex and dynamic meshwork of proteins that provides biochemical and mechanical signals governing cellular phenotypes ^20,21^. The ECM plays key roles in promoting tumor metastasis ^22^, including in HGSOC ^15,23^. It can also guide metastatic tropism, *i.e.*, the preferential dissemination or outgrowth of secondary tumors in specific organs as originally proposed by Stephen Paget in 1889 when he formulated the “seed and soil” theory ^24–27^.

Several studies have demonstrated that changes in the omental ECM occur as ovarian cancer metastases grow in this tissue ^28–30^ and *in-vitro* studies have identified roles for ECM proteins in tumor cell adhesion during their spread to peritoneal tissues ^31–33^. However, much less is known about the ECM of the mesentery compared to the omentum. We hypothesized that there might be differences in ECM composition, abundance, or organization between the greater omentum and small bowel mesentery that could explain the preferential dissemination of HGSOC to the omentum. To examine this hypothesis, we combined histological analysis and ECM-focused proteomics to characterize the ECMs of donor-matched omentum and mesentery samples. Histological analysis revealed a higher fibrillar collagen content in the mesothelial layer of the omentum as compared to the mesentery. Using a stringent quantitative proteomic pipeline, we found that the ECM composition – or matrisome – of the omentum and mesentery is highly similar, with >90% of the proteins shared between the two tissues and no significant quantitative differences among the proteins represented in both matrisomes. However, our analysis also revealed small subsets of tissue-specific ECM components, which we further validated by histological analyses when reliable antibodies were available. Altogether, our data show that the human omentum and mesentery have a highly similar matrisome and that there are thus likely additional factors explaining the preferential dissemination of HGSOC metastasis to the omentum. Last, our dataset, shared publicly with ample metadata to facilitate reuse, can also serve as a reference ECM atlas of the human omentum and mesentery for future studies.

## MATERIALS AND METHODS

### Tissue procurement

De-identified normal omental and mesentery samples were collected from deceased females over the age of 21 during autopsy (**Supplementary Table S1**). Tissues were classified as non-human-subjects research by the University of Wisconsin-Madison internal review board. Samples were excluded if there was a known history of gynecological cancer. Samples were placed in cold PBS and transported to the laboratory, where they were cut into pieces (1-2 cm^2^) and snap frozen in liquid nitrogen for proteomic analysis or fixed in 4% paraformaldehyde for paraffin embedding and sectioning at the Translational Research Initiatives in Pathology (TRIP) laboratory at the University of Wisconsin Carbone Cancer Center.

### Proteomic analysis of human omentum and mesentery samples

#### ECM enrichment

150 to 300 mg of tissue was homogenized for 2 minutes at a power setting of 12 using a Bullet Blender and stainless-steel beads of varying diameters (Navy bead lysis kit, Next Advance, Averill Park, NY) following the manufacturer’s instructions. ECM enrichment was achieved through the sequential extraction of intracellular proteins in order of decreasing protein solubility, using a subcellular protein fractionation kit (Millipore, #2145) as previously described ^34^. The efficiency of the sequential extraction of intracellular components and concomitant ECM-protein enrichment was monitored by western blot analysis, probing for collagen I (Sigma, #AB765P, used at a 1 μg/mL concentration), GAPDH (Sigma, #MAB374, used at a 5 μg/mL concentration), and actin using a serum containing anti-actin antibodies (used at a 1/5000 dilution) kindly gifted by the Hynes lab at the Massachusetts Institute of Technology, Cambridge, MA, USA.

#### Protein sample preparation for mass spectrometry

The ECM-enriched protein samples were subsequently solubilized and digested into peptides as previously described ^34^. In brief, proteins were solubilized in an 8M urea solution prepared in 100mM NH_4_HCO_3_, and protein disulfide bonds were reduced using 10mM dithiothreitol. Reduced disulfide bonds were then alkylated with 25mM iodoacetamide for 30 minutes at room temperature in the dark. The urea concentration was brought to 2M, and proteins were then deglycosylated with PNGaseF (New England Biolabs, #P0704L) for 2 hours at 37°C and digested sequentially, first with Lys-C (Thermo Scientific, #90307) for 2 hours at 37°C, and then with trypsin (Thermo Scientific, #90058), overnight at 37°C. A fresh aliquot of trypsin was added the following day, and samples were incubated for another 2 hours at 37°C. All incubations were performed under agitation. The samples were acidified with 50% trifluoroacetic acid (TFA) and desalted using Peptide Desalting Spin Columns (Pierce #89852). Peptides were reconstituted in a buffer containing 95% HPLC-grade water, 5% acetonitrile, and 0.1% formic acid, and the concentration of the peptide solution was measured using the Quantitative Colorimetric Peptide Assay kit (Pierce, #23275).

#### Peptide analysis by LC-MS/MS

Approximately 1μg of desalted peptides of each sample was analyzed at the UIC Mass Spectrometry Core facility on a quadrupole Orbitrap mass spectrometer Q Exactive HF mass spectrometer coupled to an UltiMate 3000 RSLC nano system with a Nanospray Frex Ion Source (Thermo Fisher Scientific). In brief, samples were loaded into a PepMap C18 cartridge (0.3 x 5mm, 5μm particle) trap column and then a 75 μm x 150 mm Waters BEH C18 (130A, 1.7μm, 75μm x 15cm) and separated at a flow rate of 300 nL/min. Solvent A was 0.1% formic acid (FA) in water and solvent B was 0.1% FA, 80% acetonitrile (ACN) in water. The solvent gradient of LC was 5% B in 0-3 min, 8% B in 3.5min, 8-40% B in 85 min, 40-95% B in 90min, wash 95% in 95 min, followed by 5% B equilibration until 105 min. Full MS scans were acquired in the Q-Exactive mass spectrometer over 350-1400 m/z range with a resolution of 120,000 from 5 min to 105 min. The AGC target value was 3.00E+06 for the full scan. The 20 most intense peaks with charge states 2, 3, 4, 5 were fragmented in the HCD collision cell with a normalized collision energy of 28%; these peaks were then excluded for 45s within a mass window of 1.2 m/z. A tandem mass spectrum was acquired in the mass analyzer with a resolution of 30,000. The AGC target value was 1.00E+05. The ion selection threshold was 2.00E+04 counts, and the maximum allowed ion injection time was 50 ms for full scans and 120 ms for fragment ion scans.

#### Database searching and criteria for protein identification

Spectra were searched against the UniProt human database using MaXQuant (version 2.0.3.0) with the following parameters: parent mass tolerance of 20 ppm, fragment ion mass tolerance of 20ppm, and constant modification on cysteine alkylation, variable modifications on methionine oxidation, deamidation of asparagine, glutamine, N-terminal glutamine to pyroglutamate, and hydroxylation of lysine and proline, the latter two being characteristic post-translational modifications of ECM proteins, in particular collagens and collagen-domain-containing proteins, as we previously reported ^30^. Search results were entered into Scaffold Q+S software (v5.2.0, Proteome Software, Portland, OR) for compilation, normalization, and comparison of total precursor ion intensities. Peptide identifications were accepted if they could be established with a false-discovery rate (FDR) less than 1.0% by the Scaffold Local FDR algorithm. Protein identifications were accepted if they could be established with a false-discovery rate (FDR) less than 1.0% and were identified with at least 2 distinct peptides. Proteins that contained similar peptides and could not be differentiated based on MS/MS analysis alone were grouped to satisfy the principles of parsimony. Proteins sharing significant peptide evidence were grouped into clusters. Mass spectrometry output was further annotated to identify ECM and non-ECM components. Specifically, matrisome components are classified as core-matrisome or matrisome-associated components and further categorized into groups based on structural or functional features: ECM glycoproteins, collagens, or proteoglycans for core matrisome components; and ECM-affiliated proteins, ECM regulators, or secreted factors for matrisome-associated components (**Supplemental Table S2**).

Raw mass spectrometry data and accompanying metadata file in the Sample and Data Relationship Format (SDRF) using lesSDRF ^35^ have been deposited to the ProteomeXchange Consortium ^36^ via the PRIDE partner repository ^37^ with the dataset identifier PXD061586.

### Histochemical staining

Tissue sections were deparaffinized and rehydrated in a graded ethanol series according to standard procedures.

#### Picrosirius red staining

Deparaffinized sections were incubated with picrosirius red (PSR) solution at room temperature for 1 hour. The PSR solution was prepared by dissolving 0.5 g of Direct Red 80 (Sigma-Aldrich) in 500 mL of saturated picric acid (Sigma-Aldrich). Subsequently, sections were washed with 0.5% (v/v) acetic acid in water, dehydrated in 100% ethanol, cleared in xylene, and mounted with Richard-Allan toluene-based mounting medium (Thermo Fisher Scientific). The stained sections were imaged on a Zeiss Axio Observer.Z1 inverted microscope with a Plan-Apochromat 20x/0.8 Ph2 M27 objective using the filter set for mCherry (Zeiss, Oberkochen, Germany). For each tissue, five fields of view were captured at the mesothelial edge of the tissue. The width of the ECM band from the mesothelial layer to the first adipocytes was measured in ImageJ ^38^, and the mean fluorescence intensity (MFI) of the mesothelial layer and the first row of adipocytes was measured. An additional five fields of view were captured in the adipocyte-rich bulk, and the MFI was measured using ImageJ. Measurements were averaged across the five fields of view for each patient. As we used patient-matched tissues throughout, samples were compared by a paired T-test; p<0.05 was considered statistically significant.

#### Immunohistochemical staining

Immunohistochemical staining of paired human omentum and mesentery sections (N=3) was performed at the UIC Research Histology and Tissue Imaging Core. The following antibodies were used: anti-COL12A1 antibody (Abcam #ab196619; used at a final concentration of 10μg/mL), anti-TNC antibody (Abcam #ab108930; used at a final concentration of 10μg/mL, after heat-induced epitope retrieval performed using a Tris solution at pH 9). Staining was performed with the BOND Polymer Refine Detection Kit (Leica, #DS9800) on the BOND RX automated stainer (Leica Biosystems) according to a standard preset protocol. In brief, endogenous peroxidase activity and non-specific binding sites were blocked by sequentially treating samples with peroxidase block (BOND Polymer Refine Detection Kit) and protein block (Background Sniper, Biocare Medical, #BS966) for 15 min. at room temperature. Sections were then incubated with the primary antibody for 30 min at room temperature. After several washes, signal detection was performed with a rabbit anti-HRP antibody and DAB (BOND Polymer Refine Kit) by incubating the sections for 15 min. and 10 min. at room temperature, respectively. Sections were counterstained with hematoxylin. All slides were dehydrated in an Autostainer XL and mounted with Micromount (Leica Biosystems).

Stained sections were scanned at a resolution of 40X using a Leica Aperio T2 slide scanner. Images were manually annotated using the Aperio ImageScope version 12.4.6.5003 to mark tissue boundaries and exclude artefacts such as tissue folding. The annotations were imported to Halo 3.6.4134 (Indica Labs) and analyzed by an experimenter blinded to the proteomic results using the area quantification 2.4.3 algorithm to quantify collagen XII-positive or tenascin C-positive areas across the entire tissue sections. In brief, the two-color channels were manually assigned from single-stained regions. The threshold for the two primary colors was manually determined for each staining condition. Next, two phenotypes were defined: collagen XII-positive or tenascin C-positive regions where the staining (brown color) is positive, but the counterstain (blue) is negative (phenotype 1), and a nucleus region where the counterstain is positive (phenotype 2). The positive area is calculated by dividing the area of phenotype 1 by the total tissue area.

### Statistical analysis and data visualization

Statistical analysis and data visualization were performed using Prism (GraphPad). Proportional Venn diagrams were generated with the BioVenn application: https://www.biovenn.nl/ ^39^. UpSet plots were generated using UpSetR ^40^.

## RESULTS AND DISCUSSION

### Histological characterization of the ECM of human mesentery and omentum samples

The extracellular matrix is found in distinct locations within the mesentery and omentum. An interstitial-type ECM ^20^ is found on the basal side of the mesothelial layer, referred to as the tissue edge. In contrast, a basement-membrane-type ECM ^20,41^ is found surrounding the vascular endothelium and around adipocytes (**Figure 1A**). One challenge in comparing the ECM of two tissues is the potential for variations within the tissues due to the underlying health of the donor, as conditions such as obesity result in inflammation and fibrosis of adipose tissues ^42^. Therefore, we opted to obtain matched samples of the omentum and mesentery. However, the omentum and mesentery are not commonly biopsied from patients with benign conditions, and the need for non-fixed samples for proteomic analysis prevented us from using archived samples. Therefore, we collected tissue from female decedents. To assess the overall distribution of the ECM in human mesentery and omentum, we stained tissue sections with picrosirius red, a dye that binds specifically to fibrillar collagens (primarily collagens I and III) (**Figure 1B**). We found that the overall thickness of the collagen band between the adipose and the mesothelial layer was similar between the two tissues (**Figure 1C, left panel**). While the ECM around the adipocytes is often described as a basement-membrane type ECM (lacking fibrillar collagen and containing the network-forming collagen IV), we noted in these samples the presence of fibrillar collagen between adipocytes (**Supplementary Figure S1**). We next quantified the mean fluorescence intensity corresponding to fibrillar collagen staining and observed a significantly higher collagen intensity for the region along the tissue edge relative to the adipose-rich interior region, but only for the omentum (**Figure 1C, right panel**).

**Figure 1.**
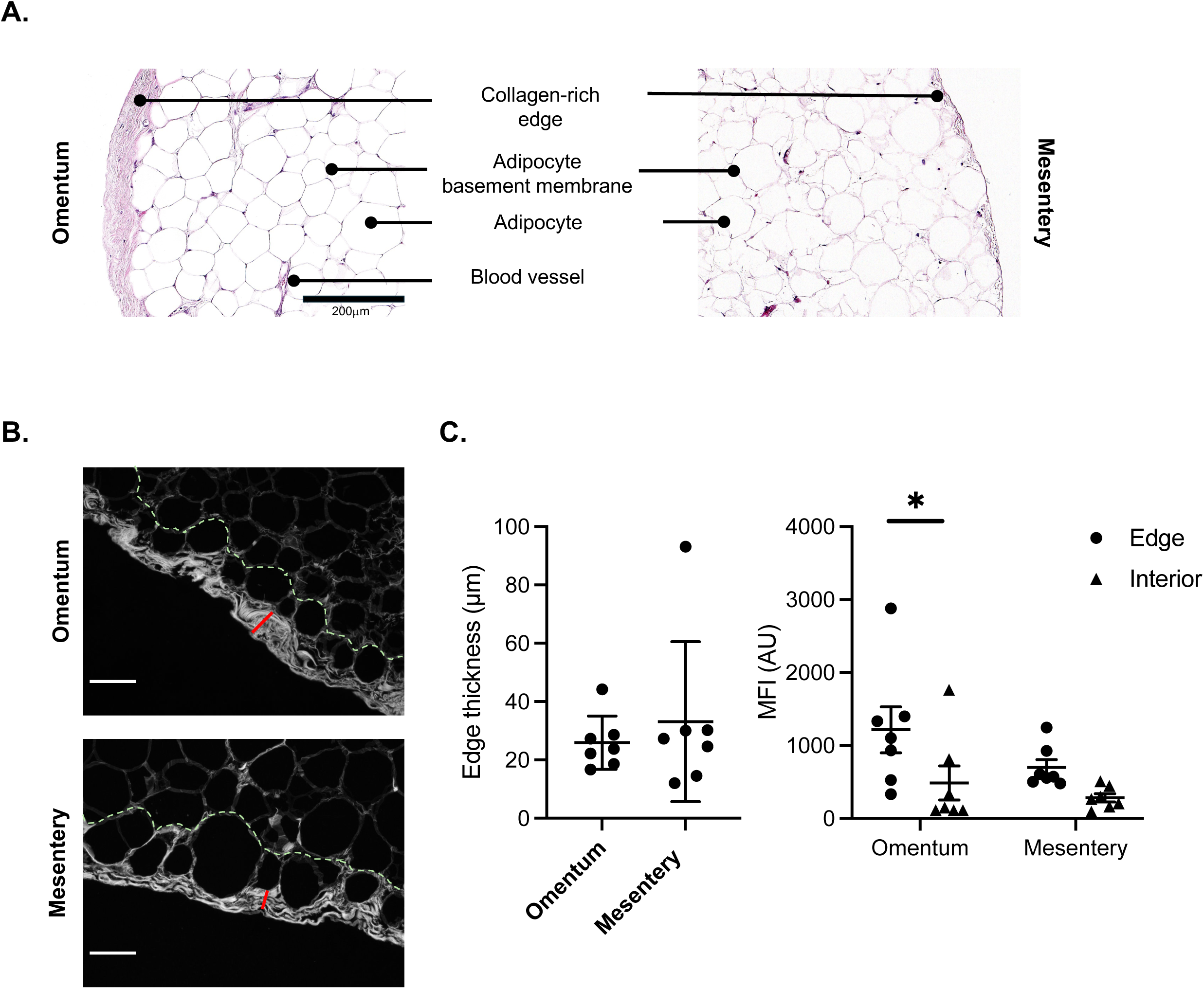
Histological characterization of the ECM of the human omentum and mesentery. **A.** Annotated hematoxylin and eosin staining of a histological section of the human omentum (left) and mesentery (right). Scale bar: 200μm. **B.** Picrosirius-red-stained images of the omentum and mesentery were analyzed for the thickness of the collagen band underlying the mesothelial layer (red line; edge) and the mean fluorescent intensity of the collagen located within the edge and first layer of adipocytes (green dotted line; interior). Scale bar: 50μm. **C.** Bar graphs represent the thickness of the tissue edge (left) and the mean fluorescence intensity of collagens in the edge (circle) or interior (triangle) region. Mean value and standard error of the mean are depicted, in addition to each data point (N=7). T-test *: p<0.05. *See also Supplementary Figure 1*.

### Proteomic pipeline to characterize the ECM composition of human mesentery and omentum samples

To characterize the molecular composition of the human mesenteric and omental ECMs, we obtained paired healthy omentum and mesentery samples from four donors (**Figure 2A**) and submitted these samples to a five-step decellularization process, extracting sequentially proteins of high to intermediate solubility, and resulting in the enrichment of highly insoluble ECM proteins ^34^> This approach is necessary to specifically cpature those proteins that are, in fact, incorporated into the ECM scaffold. We evaluated the efficiency of the decellularization using western blotting and found that intracellular proteins were completely (GAPDH) or at least partially (actin) extracted in the intermediate fractions generated through the decellularization protocol. Importantly, we observed that the ECM protein marker, collagen I, was not extracted during the procedure and was retained in the ECM-enriched fractions for both mesentery and omentum samples (**Figures 2B and 2C**).

**Figure 2.**
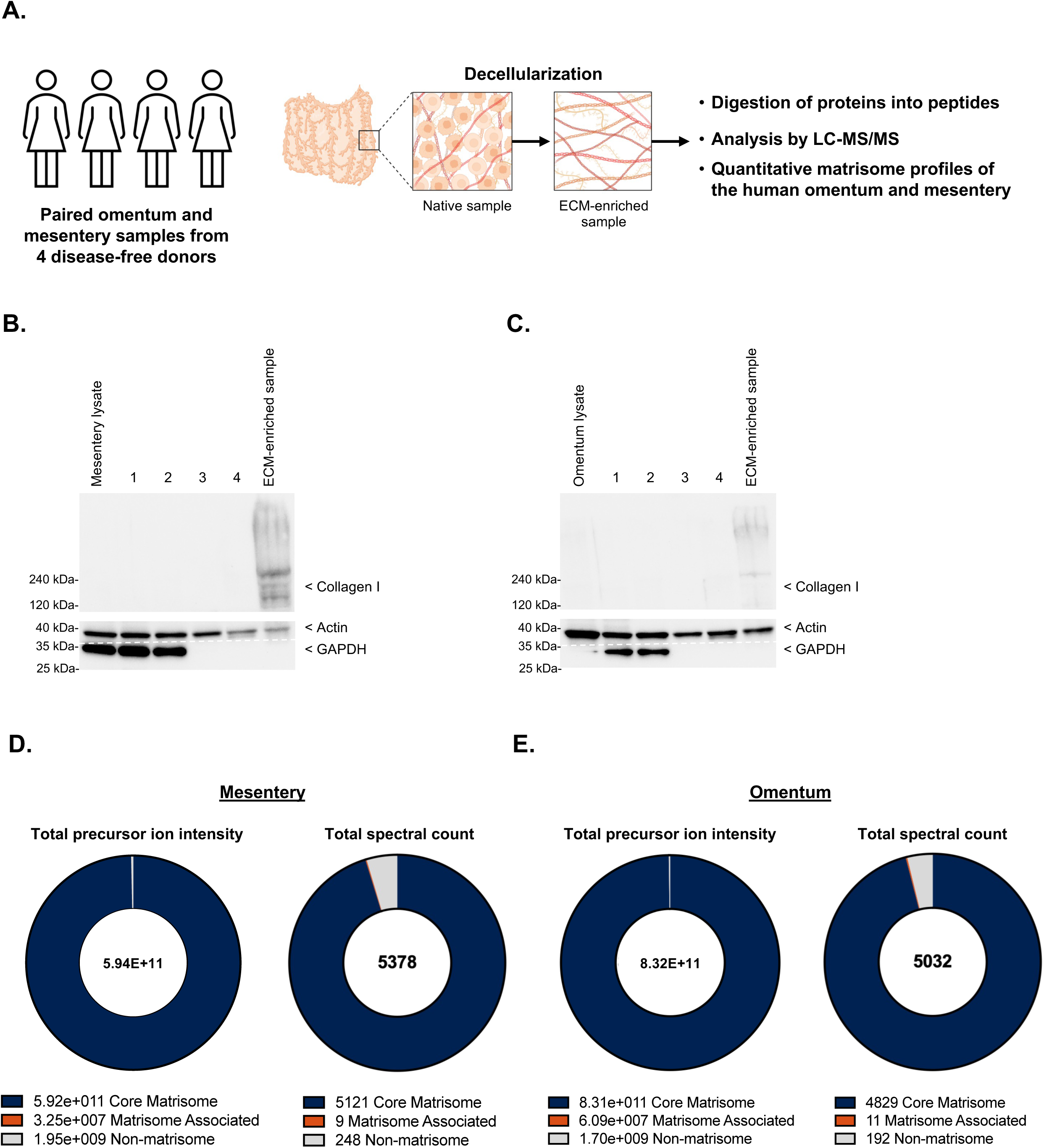
Experimental workflow to characterize the ECM of the human omentum and mesentery. **A.** Schematic representation of the experimental design of this study, created using BioRender. **B - C.** Western blots showing the sequential depletion of intracellular components (actin, GAPDH) during the 4-step decellularization process of human mesentery (**B**) and omentum (**C**) samples (fractions 1 to 4) and the resulting enrichment of ECM proteins such as collagen I. **D - E.** Donut charts represent the proportion of the total precursor ion intensity (*left*) and total spectral count (*right*) of the core matrisome (dark blue), matrisome-associated (orange), and non-matrisome (grey) components in a representative set of matched human mesentery (**D**) and omentum (**E**) ECM-enriched protein samples. *See also Supplementary Table 2*.

Next, we submitted these ECM-enriched samples to proteolytic digestion to generate peptides for proteomic analysis using liquid chromatography coupled with tandem mass spectrometry (LC-MS/MS). We analyzed the LC-MS/MS output with stringent criteria, accepting peptide and protein identifications if they could be established with a false-discovery rate (FDR) of less than 1.0% and requiring the detection of at least 2 unique peptides to further accept protein identification. We first sought to assess the efficiency of the ECM enrichment strategy. To do so, we determined the proportion of the precursor ion intensities and spectral counts attributable to matrisome vs non-matrisome proteins. These two metrics serve as proxies for protein abundance. Our results confirmed that we achieved a significant enrichment of ECM components for both mesentery and omentum samples, as the majority of the total precursor ion intensities (>99%; **Figures 2D and 2E, left panels**) and spectral count (> 95%; **Figures 2D and 2E, right panels**), were attributable to core matrisome proteins, *i.e.*, proteins forming the structure of the ECM scaffold (*see also* **Supplementary Tables S2A and B**).

### Proteomic profiling of the ECM of human mesentery samples

We next compared the proteomic profile of each of the four mesentery samples and found a substantial overlap with 22 proteins detected in all four samples, and another 27 detected in at least two independent samples (**Figure 3A** and **Supplementary Figure S2A)**. We defined the “matrisome” of the human mesentery as the ensemble of 49 proteins detected in at least two mesentery samples. The mesentery matrisome is composed of 42 core matrisome proteins, including 21 ECM glycoproteins, 16 collagens, and five proteoglycans (**Figure 3B**, **Table 1, Supplementary Table S2C**). Our analysis identified distinct protein chains of functional ECM multimers. For example, we detected the α4, α5, β1, β2, and γ1 laminin chains, encoded by the *LAMA4*, *LAMA5*, *LAMB1*, *LAMB2*, and *LAMC1* genes, respectively. Laminin chains are known to assemble selectively to form functional trimers ^43^. Our results point to the presence of four functional assemblies, the α4β1γ1, α4β2γ1, α5β1γ1, and α5β2γ1 trimers, in the basement membrane compartment of the mesentery.

**Figure 3.**
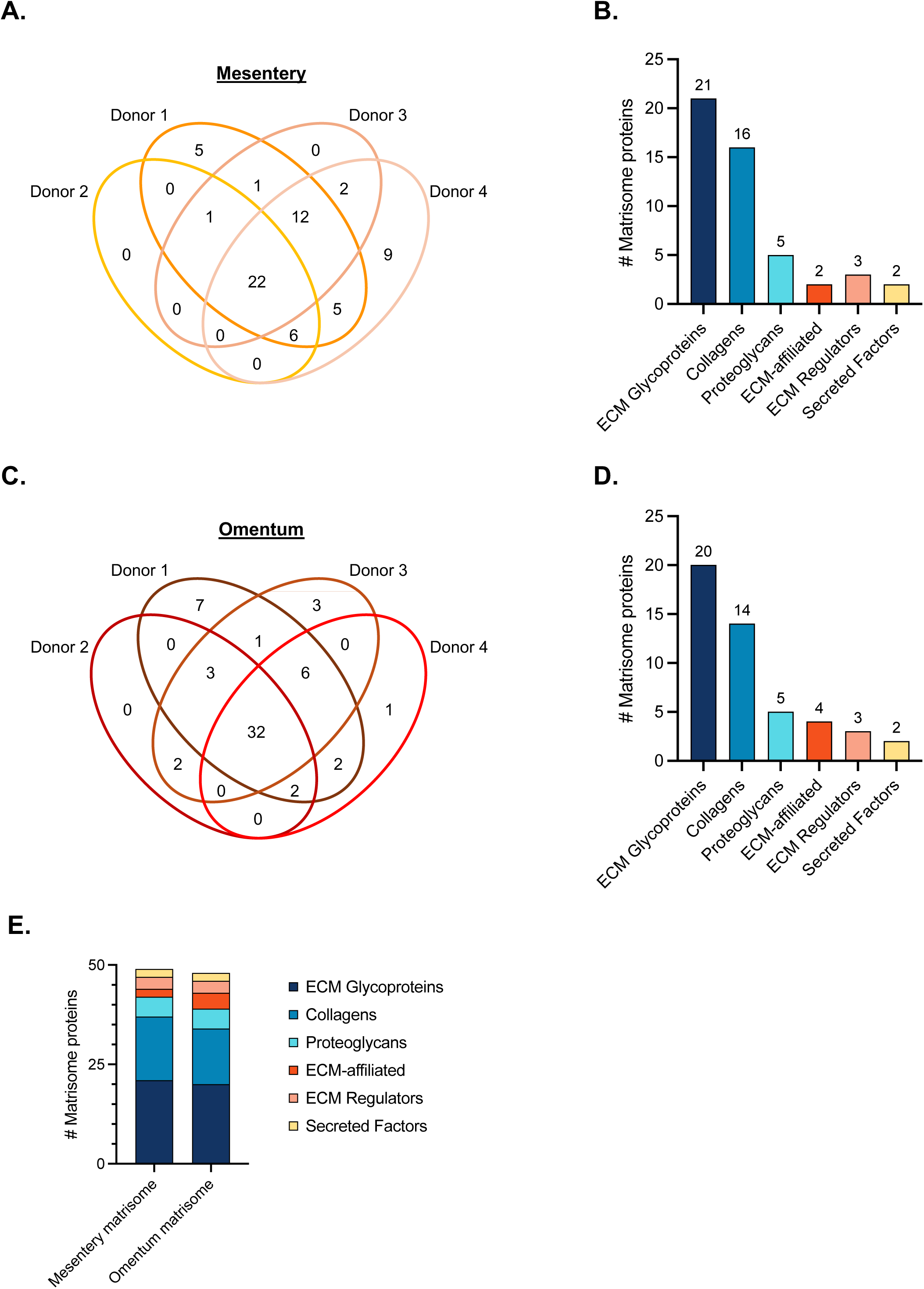
Proteomic profiling of the ECM of human mesentery and omentum samples. **A.** Venn diagram represents the number of matrisome proteins detected in each of the four mesentery samples profiled. **B.** Bar graph shows the distribution of matrisome proteins identified in at least two mesentery samples across the different matrisome categories. This ensemble of 49 proteins constitutes the mesentery matrisome. **C.** Venn diagram represents the number of matrisome proteins detected in the omentum samples for each donor. **D.** Bar graph shows the distribution of matrisome proteins identified in at least two omentum samples across the different matrisome categories. This ensemble of 48 proteins constitutes the omentum matrisome. **E.** Bar graph shows the distribution of proteins of the human mesentery (left) and omentum (right) matrisomes across the different categories of matrisome components.

**TABLE 1.** The human mesentery matrisome. The human mesentery matrisome is composed of 49 proteins identified in at least two of the four mesentery samples profiled. *Related to Figures 3A-B*

| Core matrisome |  |  | Matrisome-associated |  |  |
| --- | --- | --- | --- | --- | --- |
| ECM Glycoproteins | Collagens | Proteoglycans | ECM-affiliated | ECM Regulators | Secreted Factors |
| EFEMP1 | COL1A1 | ASPN | ANXA1 | AMBP | ANGPTL2 |
| ELN | COL1A2 | BGN | LGALS1 | LOXL1 | S100A10 |
| EMILIN1 | COL2A1 | DCN |  | TGM2 |  |
| FBLN2 | COL3A1 | HSPG2 |  |  |  |
| FBN1 | COL4A1 | LUM |  |  |  |
| FGA | COL4A2 |  |  |  |  |
| FGB | COL4A5 |  |  |  |  |
| FGG | COL5A1 |  |  |  |  |
| FN1 | COL5A2 |  |  |  |  |
| IGFBP7 | COL5A3 |  |  |  |  |
| LAMA4 | COL6A1 |  |  |  |  |
| LAMA5 | COL6A2 |  |  |  |  |
| LAMB1 | COL6A3 |  |  |  |  |
| LAMB2 | COL8A1 |  |  |  |  |
| LAMC1 | COL12A1 |  |  |  |  |
| NID1 | COL14A1 |  |  |  |  |
| NID2 |  |  |  |  |  |
| POSTN |  |  |  |  |  |
| TNC |  |  |  |  |  |
| VTN |  |  |  |  |  |
| VWF |  |  |  |  |  |

Interestingly, we identified two collagen IV chains, α1 (COL4A1) and α2 (COL4A2), that assemble to form the functional ([α1]_2_;α2) collagen IV trimer found in basement membranes. We also detected the α5 chain of collagen IV (COL4A5), which has been reported to assemble with either the α3 and α4 chains of collagen IV to form the function (α3;α4;α5) trimer or with the two α6 chains of collagen IV to form the ([α5]_2_;α6) ^41,44^. However, neither the α3, α4, or α6 chains were detected here. It is unlikely that this is due to a lack of detection due to insufficient abundance, since these proteins assemble with well-characterized 1:1:1 or 2:1 stoichiometries. It could be due to a limit in the sensitivity of our analytical pipeline that is unable to distinguish confidently highly homologous proteins with the stringent criteria imposed, or, last, it is possible that the α5 chain of collagen IV engages with other partners not identified before.

In addition to these basement membrane components, we identified collagen components of the interstitial ECM, such as the three collagen VI chains, α1 (COL6A1), α2 (COL6A2), and α3 (COL6A3), known to assemble to form the functional triple helical collagen VI ^44,45^. Collagen VI has been previously identified in omental and mesenteric metastases and correlates with a poor prognosis at the transcriptional level ^46^. While prior studies have focused on the impact of collagen VI produced by tumor cells ^46–48^, these findings support that this protein may be involved in early metastasis and become dysregulated in the tumor microenvironment. We also identified the two collagen I chains, α1 (COL1A1) and α2 (COL1A2), that must assemble to form a functional collagen I trimer ([α1]_2_;α2) ^44^. The mesentery core matrisome also includes fibrillar glycoproteins like fibronectin (FN1), fibrillin 1 (FBN1), proteins forming elastic fibers such as elastin (ELN) and the elastic microfibril interface located protein-1 (EMILIN1), the matricellular proteins ^49,50^ fibulins 2 and 3 (FBLN2 and EFEMP1), periostin (POSTN) and tenascin-C (TNC), and ECM proteins associated with hemostasis ^51^ like the three fibrinogen chains (FGA, FGB, FGG) and the von Willebrand factor (VWF).

The mesentery matrisome also included seven matrisome-associated proteins, including two ECM-affiliated proteins, three ECM regulators (including two ECM crosslinking enzymes, LOXL1 and TGM2), and two secreted factors (**Figure 3B**, **Table 1, Supplementary Table S2C**). Of note, this study provides the first proteomic characterization of the ECM of the human mesentery of disease-free individuals and can serve as the baseline for future studies focused on this tissue.

### Proteomic profiling of the ECM of human omental samples

We next compared the proteomic profile of each of the four omentum samples and found a very large overlap with 32 proteins detected in all four samples, and another 16 detected in at least two independent samples (**Figure 3C** and **Supplementary Figure S2B)**. Therefore, we define the “matrisome” of the human omentum as the ensemble of 48 proteins detected in at least two omentum samples. The omentum matrisome is composed of 39 core matrisome proteins, including 20 ECM glycoproteins, 14 collagens, and five proteoglycans (**Figure 3D**, **Table 2, Supplementary Table S2D**), a distribution similar to what we observed for the mesentery (**Figure 3E**). Our analysis identified the same laminin and collagen chains and trimer as those detected in the mesentery. The omentum matrisome also included nine matrisome-associated proteins, including four ECM-affiliated proteins, three ECM regulators (including two ECM crosslinking enzymes, LOXL1 and TGM2, also found in the mesentery matrisome), and two secreted factors (**Figure 3D**, **Table 2, Supplementary Table S2D**). While there is no prior study on the ECM composition of the human mesentery, we and others have reported the composition of the ECM of the human omentum, and these studies have typically identified a larger number of proteins ^28,30,52^. This can be explained by the fact that we focused here on the identification of proteins of the insoluble ECM scaffold after sample decellularization, as opposed to previous studies that have performed proteomic analyses on the bulk omental samples or urea-soluble omental samples. We have also applied stringent criteria to accept protein identification, including strict false-discovery rate thresholds and strict cohort-level cut-offs (*see Materials and Methods*). Last, since we focused on the most insoluble ECM protein fraction, fibrillar collagens that contribute hundreds of spectra, tend to saturate the MS signal, decreasing our ability to identify proteins present in lower abundance. To alleviate this, we attempted to fractionate samples using basic reverse-phase liquid chromatography, a method commonly employed to achieve greater depth in proteomic studies; however, because of the dominance of collagen-derived peptides, this approach fell short.

**TABLE 2.**
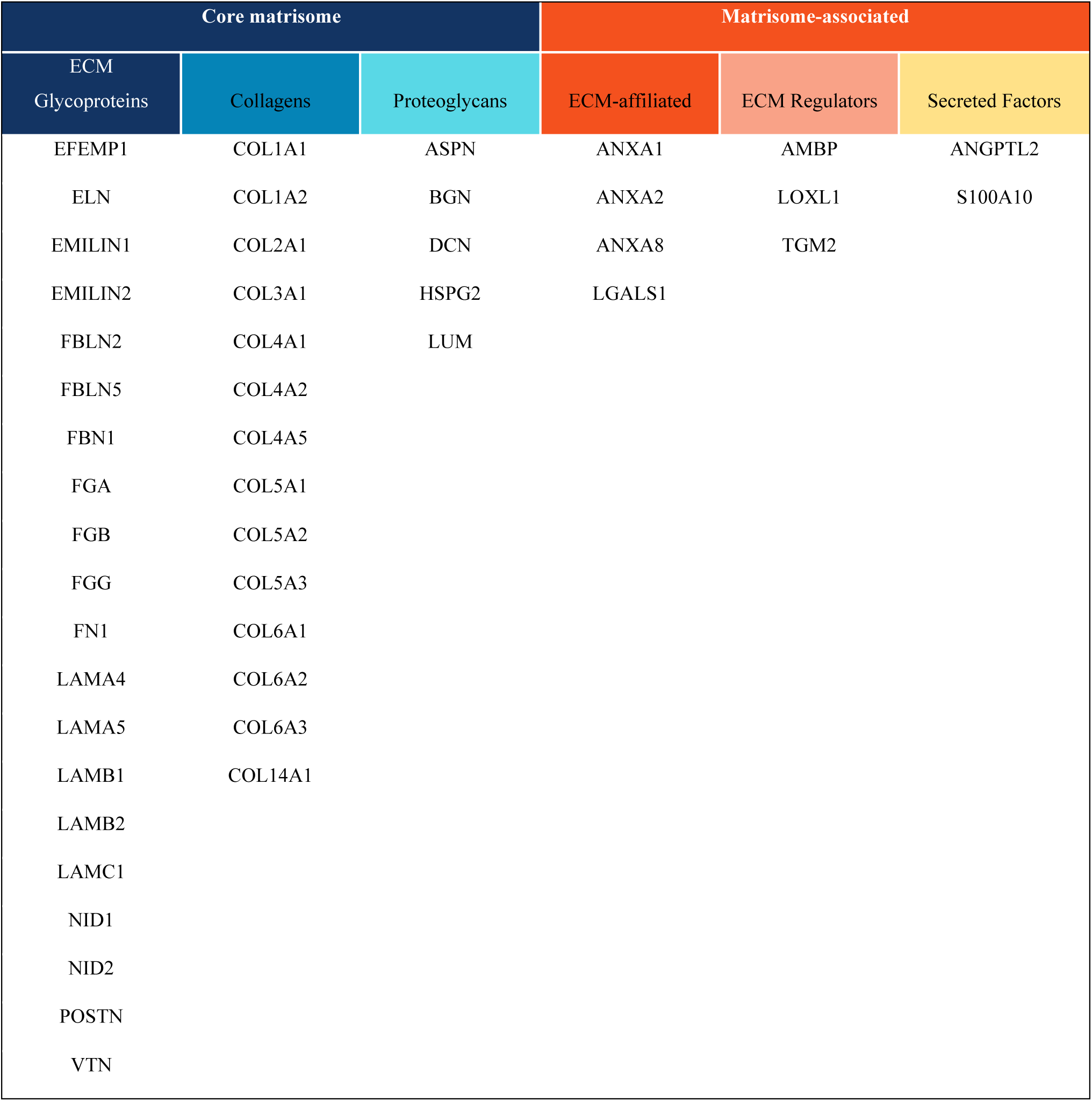
The human omentum matrisome. The human omentum matrisome is composed of 48 matrisome proteins identified in at least two of the four omentum samples profiled. *Related to Figures 3C-D*

| Core matrisome |  |  | Matrisome-associated |  |  |
| --- | --- | --- | --- | --- | --- |
| ECM Glycoproteins | Collagens | Proteoglycans | ECM-affiliated | ECM Regulators | Secreted Factors |
| EFEMP1 | COL1A1 | ASPN | ANXA1 | AMBP | ANGPTL2 |
| ELN | COL1A2 | BGN | ANXA2 | LOXL1 | S100A10 |
| EMILIN1 | COL2A1 | DCN | ANXA8 | TGM2 |  |
| EMILIN2 | COL3A1 | HSPG2 | LGALS1 |  |  |
| FBLN2 | COL4A1 | LUM |  |  |  |
| FBLN5 | COL4A2 |  |  |  |  |
| FBN1 | COL4A5 |  |  |  |  |
| FGA | COL5A1 |  |  |  |  |
| FGB | COL5A2 |  |  |  |  |
| FGG | COL5A3 |  |  |  |  |
| FN1 | COL6A1 |  |  |  |  |
| LAMA4 | COL6A2 |  |  |  |  |
| LAMA5 | COL6A3 |  |  |  |  |
| LAMB1 | COL14A1 |  |  |  |  |
| LAMB2 |  |  |  |  |  |
| LAMC1 |  |  |  |  |  |
| NID1 |  |  |  |  |  |
| NID2 |  |  |  |  |  |
| POSTN |  |  |  |  |  |
| VTN |  |  |  |  |  |

### Comparative analysis of the human omentum and mesentery matrisomes

We first compared the ECM profiles of paired omentum and mesentery samples and found a very large overlap at the individual donor level (**Supplementary Figure S2C**). Moreover, the comparative analysis of the matrisomes of the human mesentery and omentum reveals an overall very similar ECM composition, both in terms of representation of the different categories of matrisome components (**Figure 3E**) and in overall protein abundances (**Figure 4A**), with fibrillar collagens (COL1A1, COL1A2, COL3A1) contributing to the majority of the protein signal and dominating ECM composition by two orders of magnitude over the rest of the matrisome (**Figure 4A** and **Supplementary Table S2B**). This result, in combination with the PSR staining demonstrating a thicker collagen band surrounding the adipose with a thin network between individual adipocytes, suggests that fibrillar collagens play an essential role in the overall organization of these tissues. Collagen I is further increased in tissues with metastases ^29^, where it remains the dominant component ^28,30^.

**Figure 4.**
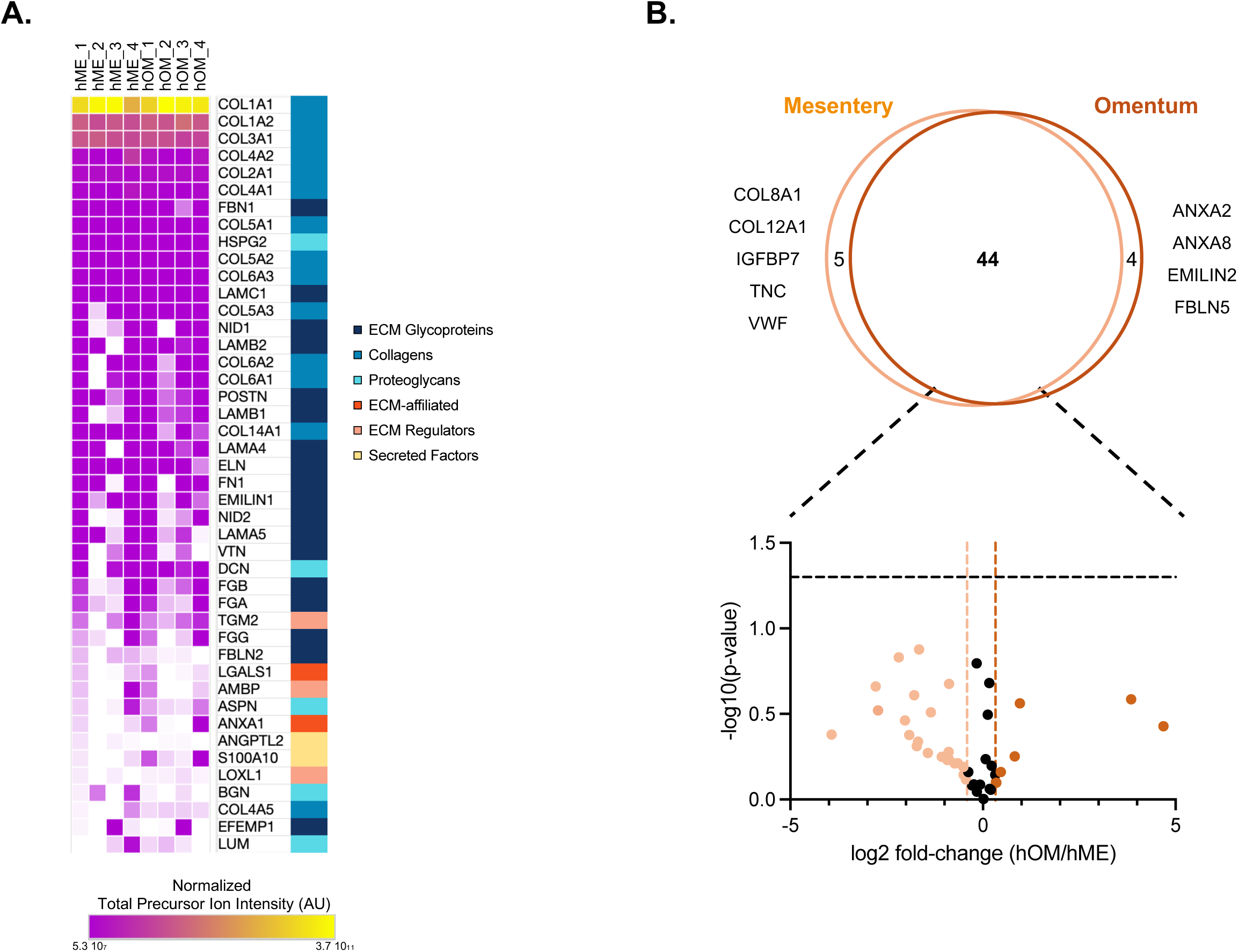
Comparison of the human omentum and mesentery matrisomes. **A.** Heatmap of normalized total precursor ion intensities of the 44 proteins detected in both human mesentery and omentum matrisomes. **B.** Proportional Venn diagram represents the overlap between the mesentery and omentum matrisomes. Volcano plot represents the fold-change average protein abundance inferred from total precursor ion intensity (*see Methods*) between the omentum and mesentery. Dotted lines along the x-axis indicate a 25% lower (left quadrant) or higher (right quadrant) protein abundance in the omentum as compared to the mesentery. The dotted line along the y-axis depicts the statistical significance cut-off set to p<0.05.

The two tissues share 44 matrisome proteins (**Figure 4B**) and notably share the same proteoglycanome composed of asporin (ASPN), biglycan (BGN), decorin (DCN), lumican (LUM), and the basement-membrane-associated proteoglycan perlecan (HSPG2). The quantitative analysis of the abundance of these 44 proteins in mesenteric and omental samples did not reveal any statistically significant differences. However, our analysis led to the identification of five matrisome proteins in the mesentery matrisome but not in the omentum matrisome (**Figure 4B**). These include the network-forming collagen VIII (COL8A1), the fibril-associated collagen with interrupted triple helix collagen XII (COL12A1), the insulin-like growth factor binding protein (IGFBP7), the matricellular protein tenascin-C (TNC) ^53^, and the von Willebrand factor (VWF). Conversely, we identified four matrisome proteins characteristic of the omentum matrisome (**Figure 4B**). These include two components of the core matrisome, the elastic microfibril interface located protein-2 (EMILIN2), and the matricellular protein fibulin 5 (FBLN5), neither of which has been clearly associated with HGSOC metastasis. We also detected two ECM-affiliated proteins, annexins A2 and A8 (ANXA2, ANXA8), in the omentum but not in the mesentery. Annexin A2 has previously been shown to be expressed in the mesothelial layer of the omentum, and weekly treatments with a blocking antibody against annexin A2 resulted in slower tumor growth in a xenograft model of SKOV3 cells ^54^.

### Collagen XII and tenascin C present tissue-specific distribution patterns

Two of the proteins detected in the mesenteric but not omental matrisome, collagen XII and tenascin C, have previously been associated with tumor metastasis and metastatic tropism across several cancer types, including breast and pancreatic cancers ^55–60^. Somewhat surprisingly, our analysis found them associated with the ECM of the mesentery, the tissue that is not the preferential site of metastatic dissemination of HGSOC. We thus sought to validate the proteomic data using immunohistochemistry. We found that the abundance of collagen XII (**Figure 5A**) and tenascin C (**Figure 5B**) supported the proteomic results, with a higher abundance in mesentery samples as compared to omental samples. Specifically, collagen XII, which is a fibril-associated collagen with interrupted triple helices ^44^, presented a fibrillar pattern between adipocytes of the mesentery characterized by short and distinct structures (**Figure 5A, left panel**), in agreement with its known association with fibrillar collagens, collagens I and III. This pattern was absent in the omentum samples analyzed. In contrast, the matricellular protein tenascin C presents a more homogeneous pattern surrounding adipocytes, suggesting a possible association with the basement membrane that continuously surrounds these cells. We also observed an abundant staining of the large vessels of the mesentery but not of the omentum in two of the three pairs of samples analyzed (**Figure 5B, left panel**). Last, it is important to note that very low levels of collagen XII and tenascin C were still detected in the omental samples (**Figures 5A and B, right panels**). This can be attributable to the fact that our proteomic pipeline focused on the insoluble ECM, while immunohistochemistry will detect proteins regardless of their solubility or level of incorporation in the ECM.

**Figure 5.**
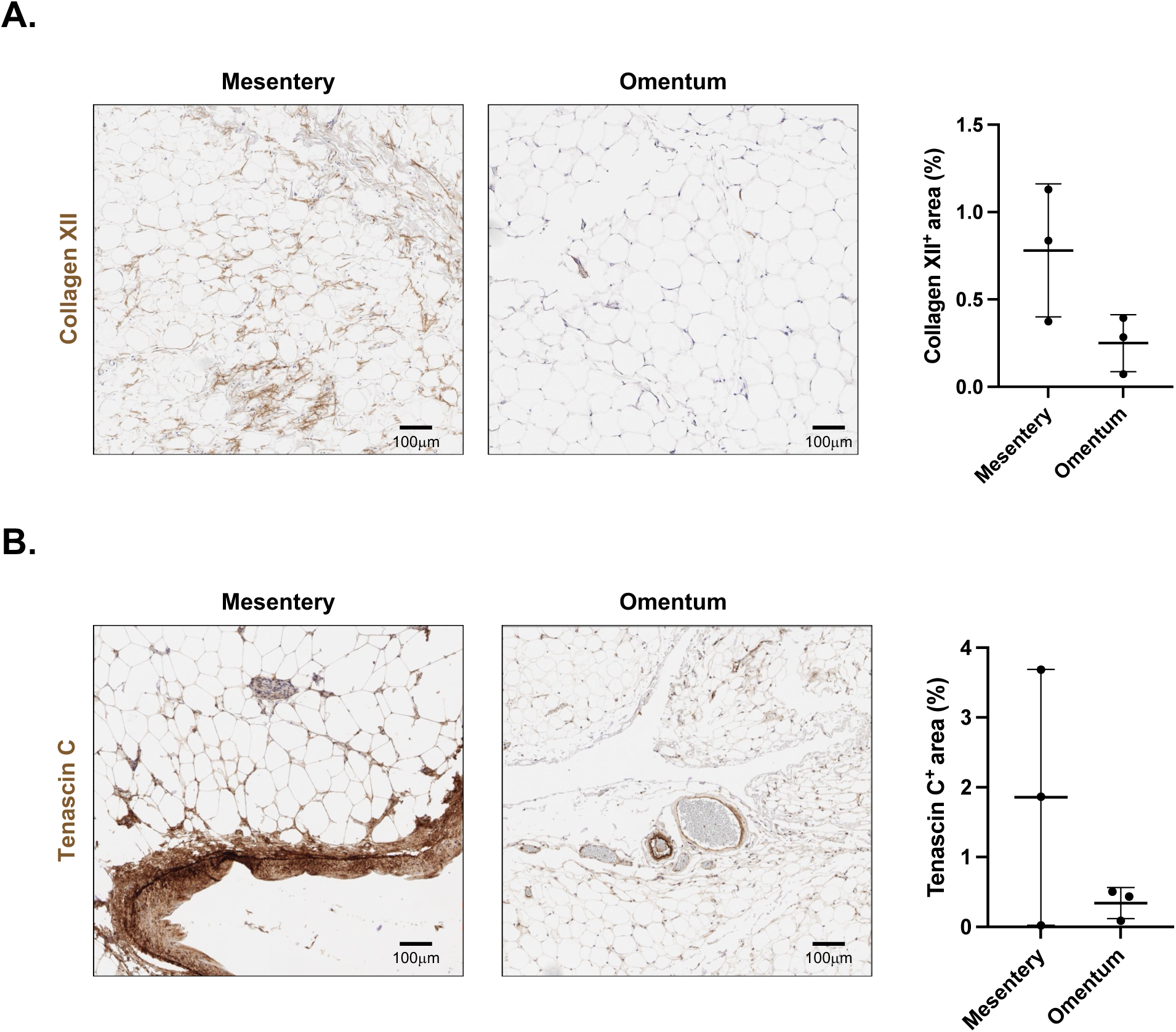
Comparison of the pattern of distribution of collagen XII and tenascin C in human mesentery and omentum samples. **A.** Representative images of 5µm-thick tissue sections stained with an anti-collagen XII antibody confirming the proteomic data showing the presence of collagen XII in the mesenteric ECM and its absence in the omental ECM. Dot plot represents the proportion of collagen XII-positive tissue area in three paired omentum and mesentery samples. Scale bar: 100μm. **B.** Representative images of 5µm-thick tissue sections stained with an anti-tenascin C antibody confirming the proteomic data showing the presence of tenascin C in the mesenteric ECM and its absence in the omental ECM. Dot plot represents the proportion of tenascin C-positive tissue area in three paired omentum and mesentery samples. Scale bar: 100μm.

## CONCLUSION

Altogether, our data show that the protein compositions of the ECM of the human omentum and mesentery are highly similar. This is consistent with recent studies that support the idea that the mesentery is a continuous organ and that the omentum develops when a ‘bubble’-like structure forms in the upper mesentery ^61^. While the similarities in the matrisome of healthy tissues do not provide an obvious explanation for the predilection for omental metastases, future work should aim to test the possible contribution of the ECM proteins presenting a tissue-specific expression to guiding the metastatic tropism of HGSOC. This can be accomplished using tissue engineering strategies that provide control over the composition of the ECM ^14^. While prior studies have focused on the impact of proteins that change drastically in abundance during metastasis, analysis of the matrisome abundance supports the use of fibrillar collagens as the base material for *in-vitro* models of the omentum and mesentery to study the initial metastasis of ovarian cancer cells to these sites ^32,33^. For example, we have recently demonstrated that the fibrillar structure of collagen I provides adhesion cues for ovarian cancer cells ^32^. In addition, it is well known that the omental matrisome changes drastically during tumor progression. Therefore, it remains possible that the tissues diverge during this process to provide distinct pre-metastatic and metastatic microenvironments.

## ACKNOWLEDGMENTS

The authors would like to thank Dr. Hui Chen and Lasanthi Jayathilaka from the UIC Mass Spectrometry Core facility and Dr. Maria Sverdlov and Dr. Ryan Easton from the UIC Research Histology and Tissue Imaging Core for their technical assistance.

## CRediT AUTHOR STATEMENT

**JC**: Investigation, Methodology, Formal Analysis, Visualization, Writing

**CG**: Investigation, Methodology, Formal Analysis

**DP:** Formal Analysis

**NY**: Investigation, Methodology, Formal Analysis

**INT**: Investigation, Methodology, Formal Analysis

**JS**: Investigation, Formal Analysis

**EGB**: Resources

**PKK**: Conceptualization, Methodology, Resources, Formal Analysis, Visualization, Writing, Supervision, Funding Acquisition, Project Administration

**AN**: Conceptualization, Methodology, Resources, Formal Analysis, Visualization, Writing, Supervision, Funding Acquisition, Project Administration

## DATA AVAILABILITY

Raw mass spectrometry data and accompanying metadata file in the Sample and Data Relationship Format (SDRF) ^35^ have been deposited to the ProteomeXchange Consortium ^36^ via the PRIDE partner repository ^37^ with the dataset identifier PXD061586. The raw data will be made publicly available upon acceptance of the manuscript.

## SUPPLEMENTARY FIGURE LEGENDS

**Supplementary Figure S1.**
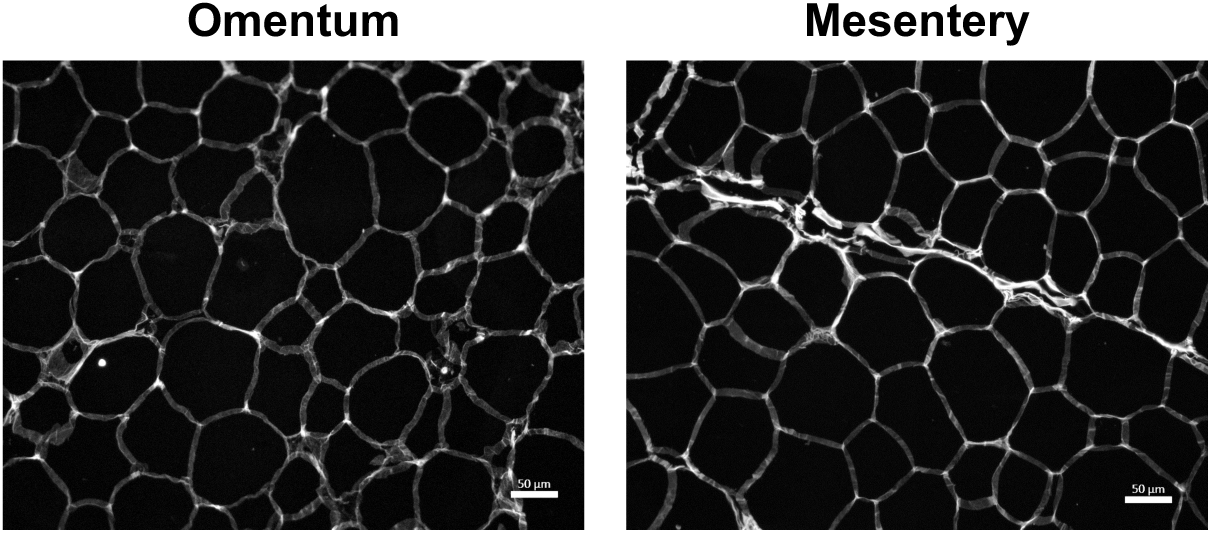
Picrosirius red staining of the adipose region of human omentum and mesentery samples. Representative images of picrosirius-red staining of the adipose regions of the omentum and mesentery used to quantify the interior collagen content in Figure 1C. Scale bar: 50µm.

**Supplementary Figure S2.**
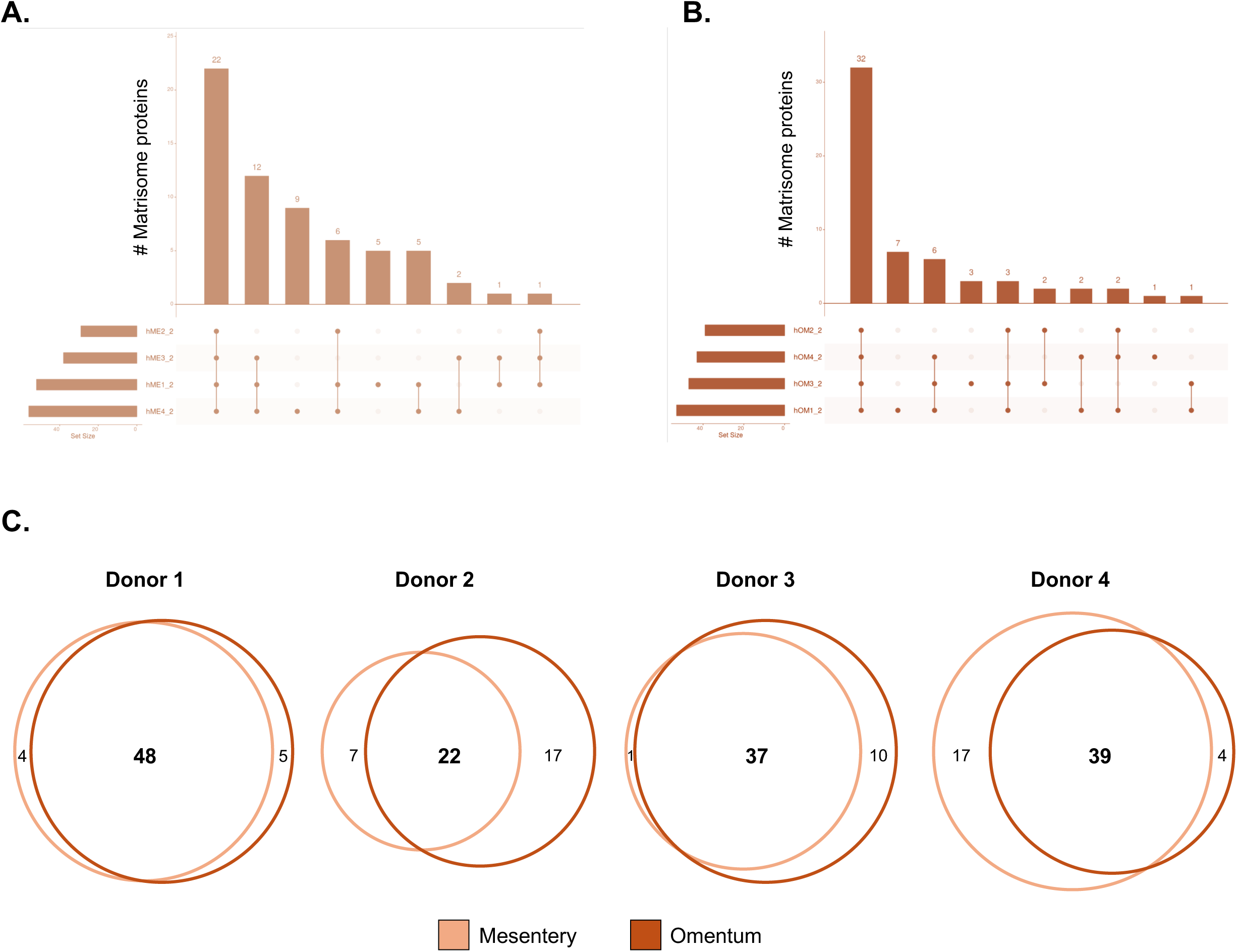
Comparison of the ECM composition of the human omentum and mesentery. **A.** UpSet plot provides a quantitative representation of the overlap between the proteins detected in each of the four mesentery samples profiled. **B.** UpSet plot provides a quantitative representation of the overlap between the proteins detected in each of the four omentum samples profiled. **C.** Proportional Venn diagrams represent the overlap between the matrisome proteins identified in paired mesentery (orange) and omentum (brown) samples from four disease-free donors.

## SUPPLEMENTARY TABLES

**Supplementary Table S1.**
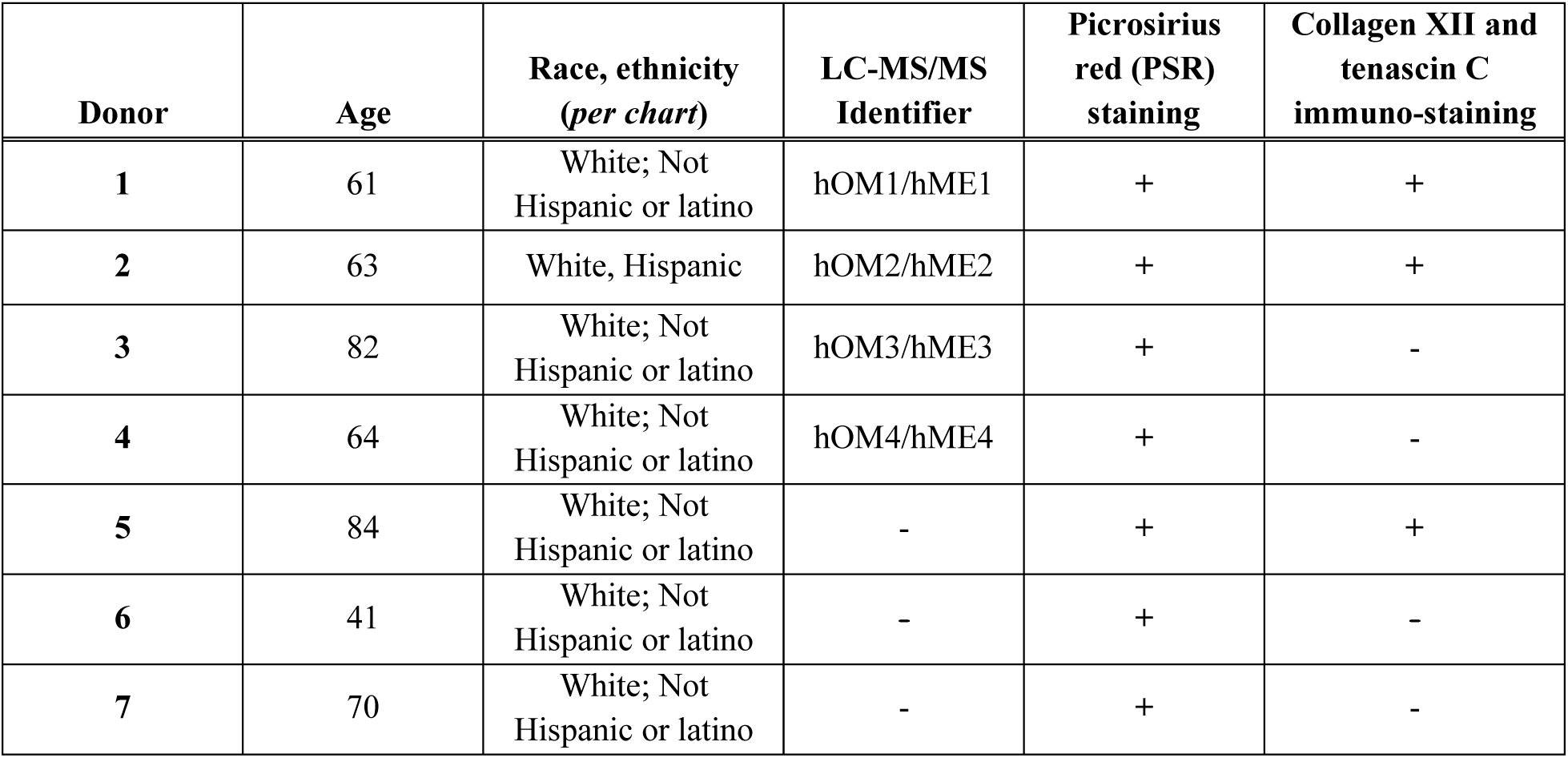
Demographic data and information on samples used for proteomic and histochemical analyses.

| **Donor** | **Age** | **Race, ethnicity**<br>**(*per chart*)** | **LC-MS/MS**<br>**Identifier** | **Picrosirius**<br>**red (PSR)**<br>**staining** | **Collagen XII and**<br>**tenascin C**<br>**immuno-staining** |
| --- | --- | --- | --- | --- | --- |
| **1** | 61 | White; Not<br>Hispanic or latino | hOM1/hME1 | + | + |
| **2** | 63 | White, Hispanic | hOM2/hME2 | + | + |
| **3** | 82 | White; Not<br>Hispanic or latino | hOM3/hME3 | + | - |
| **4** | 64 | White; Not<br>Hispanic or latino | hOM4/hME4 | + | - |
| **5** | 84 | White; Not<br>Hispanic or latino | - | + | + |
| **6** | 41 | White; Not<br>Hispanic or latino | - | + | - |
| **7** | 70 | White; Not<br>Hispanic or latino | - | + | - |

**Supplementary Table S2. Complete proteomic dataset S2A. Complete mass spectrometry dataset**

Columns G to M report the total precursor ion intensity values for each sample

Columns O to V report the total number of spectra for each sample

**S2B. List of all matrisome proteins detected**

**S2C. List of proteins composing the human mesentery matrisome**

**S2D. List of proteins composing the human omentum matrisome**

